# Metabolomics investigation of post-mortem human pericardial fluid

**DOI:** 10.1101/2023.02.28.530436

**Authors:** Alberto Chighine, Matteo Stocchero, Giulio Ferino, Fabio De-Giorgio, Celeste Conte, Matteo Nioi, Ernesto d’Aloja, Emanuela Locci

## Abstract

**Introduction:** Due to its peculiar anatomy and physiology, the pericardial fluid is a biological matrix of particular interest in the forensic field. Despite this, the available literature has mainly focused on post-mortem biochemistry and forensic toxicology, while to the best of authors’ knowledge post-mortem metabolomics has never been applied. Similarly, estimation of the time since death or Post-Mortem Interval based on pericardial fluids has still rarely been attempted.

**Objectives:** We applied a metabolomic approach based on ^1^H Nuclear Magnetic Resonance Spectroscopy to ascertain the feasibility of monitoring post-mortem metabolite changes on human pericardial fluids with the aim of building a multivariate regression model for Post-Mortem Interval estimation.

**Methods:** Pericardial fluid samples were collected in 24 consecutive judicial autopsies, in a time frame ranging from 16 to 170 hours after death. The only exclusion criterion was the quantitative and/or qualitative alteration of the sample. Two different extraction protocols were applied for low molecular weight metabolites selection, namely ultrafiltration and liquid-liquid extraction. Our metabolomic approach was based on the use of ^1^H Nuclear Magnetic Resonance and multivariate statistical data analysis.

**Results:** The pericardial fluid samples treated with the two experimental protocols did not show significant differences in the distribution of the metabolites detected. A post-mortem interval estimation model based on 18 pericardial fluid samples was validated with an independent set of 6 samples, giving a prediction error of 33 - 34 hours depending on the experimental protocol used. By narrowing the window to post-mortem intervals below 100 hours, the prediction power of the model was significantly improved with an error of 13-15 hours depending on the extraction protocol. Choline, glycine, ethanolamine, and hypoxanthine were the most relevant metabolites in the prediction model.

**Conclusion:** The present study, although preliminary, shows that PF samples collected from a real forensic scenario represent a biofluid of interest for post-mortem metabolomics, with particular regard to the estimation of the time since death.

## 1 Introduction

Pericardial fluid (PF) is a pale-yellow serous fluid present in the pericardial cavity, a virtual space that normally surrounds the heart with a volume, under physiological conditions, between approximately 15 and 60 mL. Historically it was believed that its physio-pathological role was limited to guarantee the cardiac cycle from the mechanical point of view, favouring the alternation of cardiac expansions and contractions. More recently, it has been demonstrated that PF production results not only on plasma ultrafiltration since its composition appears to be also sustained by secretion from pericardial mesothelial cells and myocardial interstitial space. Thus, this particular biofluid is supposed to carry important information on the whole pericardial and cardiac homeostasis, with growing evidence indicating several bioactive molecules to be stored in the PF, suggesting a potential paracrine activity [1].

Scientific attention to PF has been limited by obvious and shareable ethical issues, as it is not possible to collect this biofluid from healthy donors, PF collections were always performed during open-heart surgery. In this setting, PF is certainly influenced by the pathological conditions affecting the surrounding structures (*i*.*e*., valvular, or ischemic cardiopathies), for which surgery is performed [2].

Recently, the attention of the scientific community has been devoted to PF chemical and biochemical composition [3-5], but little is known about the metabolic features of this particular biofluid. To our knowledge, only two papers can be found in literature dealing with PF lipidomics [6] and metabolomics [7]. The first was conducted on an animal model (rabbit) of hypercholesterolemia and coronary atherosclerosis, whereas the second investigated PF and plasma metabolome of patients undergoing cardiac surgery for coronary artery bypass grafting. Despite its use for forensic toxicological and biochemical purposes [8-12], post-mortem human PF has never been investigated with a metabolomic approach.

Our research group is focused on the application of metabolomics to address classical forensic issues by the analysis of different biological matrices on human and animal models [13], such as the identification of specific biofluids [14], the diagnosis of perinatal asphyxia [15, 16], the PMI estimation [17-20], the differential diagnosis of asphyxial deaths [21], and the diagnosis of drug-related fatalities [22]. Death is nowadays increasingly recognized as a dynamic rather than a static phenomenon [23], and we previously observed that multivariate approaches are more suitable than univariate to describe this multiparametric phenomenon. Time since death represents indeed the main driving force of post-mortem metabolome changes and we have previously highlighted the need for PMI standardization when post-mortem metabolomics is performed for aims different than PMI [14].

PMI estimation still represents one of the biggest forensic conundrums, with a large body of forensic literature aiming at the discovery of accurate and reliable tools for its determination, sadly often far away from practical application [25]. The ideal biofluid to be studied in metabolomics for PMI estimation should be relatively stable to display monitorable time-related metabolome modifications. Such characteristics are usually found in ocular biofluids due to the unique anatomical and physiological features [26, 27]. Like ocular biofluids, PF appears suitable for post-mortem investigations as its confined anatomy and physiology may offer a relative resistance to post-mortem autolysis and putrefaction. Furthermore, this biofluid has shown remarkable stability in an animal model, with a slow turnover rate ranging between 5.4 and 7.2 hours [28-29]. To date, only one investigation aimed at estimating PMI on human PF was performed and was based on changes in electrolyte concentration over a time window of 2.5-85.0 hours [30]. Another study, performed on an animal model (rabbit), investigated shorter PMIs (up to 48 hours) based on Fourier transform infrared spectroscopy and chemometrics [31].

To the best of our knowledge, this is the first study dealing with post-mortem human PF metabolomics. The objective of this proof-of-concept study was to investigate the feasibility of PF metabolomics to estimate PMI. To this aim, PF samples collected during 24 consecutive judicial autopsies were analysed with ^1^H Nuclear Magnetic Resonance (NMR) spectroscopy following two different experimental protocols for the analysis of the metabolome.

## 2 Materials and Methods

### 2.1 Sample collection

PF samples were collected during 24 consecutive judicial autopsies performed at Forensic and Legal Medicine Institute of University of Cagliari. After routine opening of the chest wall, pericardium was anteriorly dissected through a reversal ‘Y’ incision which allowed pericardial cavity to be exposed. The cardiac apex was gently pulled up and declivous PF was collected with a sterile no-needled syringe. Collection was not performed in case of pathological pericardial effusion and/or macroscopic blood contamination. Not all the PMIs were precisely known, being estimated based on circumstantial evidence and/or classical thanatological signs. After collection, PFs were immediately stored at -80°C.

### 2.2 Sample preparation for NMR analysis

Before NMR analysis, samples were thawed, centrifuged for 10 min at 13000 g and 4°C and the supernatant was divided in two aliquots, that were treated in two different ways for protein removal. An aliquot of 500 μl was submitted to ultrafiltration (U) while the second was extracted using a liquid-liquid extraction (LLE) procedure. Ultrafiltration was performed using 10 kDa filter units (Amicon-10kDa; Merck Millipore, Darmstadt, Germany) for 25 min at 13000 g and 4°C. Filters were previously washed out from glycerol by adding 500 μl of distilled water and by centrifuging for 5 min at 13000 g at room temperature for 15 times. For the NMR analysis, 300 μl of filtered PF were diluted with 400 μl of a 0.09 M phosphate buffer solution (pH=7.4) in D_2_O (99,9%, Cambridge Isotope Laboratories Inc, Andover, USA) containing the internal standard sodium 3 (trimethylsilyl)propionate-2,2,3,3,-*d*_*4*_ (TSP, 98 atom % D, Sigma-Aldrich, Milan) at a 0.21 mM final concentration, and 650 μl of the final solution were transferred into a 5 mm NMR tube. For LLE, an aliquot of 400 μl was mixed with 1200 μl of a chloroform: methanol (1:1, both HPLC grade, Honeywell, Germany) solution and 175 μl of Milli-Q water. The samples were vortexed and centrifuged at 1700 g for 30 min at room temperature. The upper hydrophilic phase was dried overnight using a speed vacuum centrifuge (Eppendorf concentrator plus, Eppendorf AG, Hamburg, Germany). The dried samples were rehydrated by adding 300 μl of D_2_O and 400 μl of a 0.09 M phosphate buffer solution (pH=7.4) in D_2_O containing TSP at a 0.21 mM final concentration. 650 μl of the resulting solution were transferred into a 5 mm NMR tube. One PF sample was analysed without any pre-treatment. After thawing and centrifugation, 300 μl of PF were diluted with 400 μl of the phosphate buffer solution previously used, and 650 μl of the final solution were transferred into a 5 mm NMR tube.

### 2.3 ^1^H NMR experiments and data processing

^1^H NMR experiments were carried out on a Varian UNITY INOVA 500 spectrometer (Agilent Technologies, CA, USA) operating at 499.839 MHz. Spectra were acquired at 300K using a spectral width of 6000 Hz, a 90° pulse, an acquisition time of 1.5 sec, a relaxation delay of 2 sec, and 256 scans. Non treated PF and UPF spectra were acquired using the standard 1D NOESY pulse sequence for water suppression with a mixing time of 1 ms and a recycle time of 21.5 s, while in LLEPF spectra the residual water signal was suppressed by applying a standard presat pulse sequence with low-power radiofrequency irradiation for 2 sec during relaxation. Spectra were processed using MestReNova software (Version 9.0, Mestrelab Research S.L.). The free induction decays (FID) were multiplied by an exponential weighting function equivalent to a line broadening of 0.5 Hz and zero-filled to 128K before Fourier transformation. All spectra were phased, baseline corrected, and referenced to TSP at 0.00 ppm. The assignment of the metabolites in the ^1^H NMR spectra was performed based HMDB database (http://www.hmdb.ca), Chenomx NMR Suite 8.2 Library (Chenomx Inc., Edmonton, Canada), and comparison with spectra of standard compounds recorded using the same experimental conditions. Moreover, using the Chenomx NMR Suite Profiler tool a set of 50 quantified metabolites was obtained, after the exclusion of exogenous metabolites such as ethanol, caffeine and drugs (see Supplementary Table 1). The final data set was exported as a text file for multivariate statistical data analysis. Data were autoscaled prior to performing data analysis.

### 2.4 Multivariate statistical data analysis

Exploratory data analysis was performed by Principal Component Analysis (PCA) [**32**] to discover outliers and specific trends in the data. Regression models to estimate PMI from the quantified metabolites were based on orthogonally Constrained PLS2 (oCPLS2) [**33**]. The application of orthogonal constraints to PLS-regression was necessary to avoid that the factor age affected the models. Indeed, oCPLS2 guaranteed score components orthogonal to age, modelling data variation mainly associated to PMI. Relevant predictors were discovered by stability selection using the Variable Influence on Projection (VIP) score in variable selection [**34**]. Specifically, 200 subsamples were extracted by Binary Matrix Sampling with probability 0.7. For each subsample, the subset of predictors that generated the oCPLS2 model with the best performance in cross-validation was calculated. The frequency of selection of each predictor was compared with that of a random choice to discover the relevant predicators. A significance level of 0.05 was assumed. Repeated 5-fold cross-validation with 20 repetitions was applied in model optimization maximising the R^2^ on the validation set, i.e., Q^2^. Randomization test was applied to discover over-fitting and to assess the reliability of the model. The data variation of the relevant metabolites discovered by PLS-based analysis was investigated by Multiple Linear Regression (MLR) considering the factors age and PMI. A stratified random selection procedure based on the different ranges of PMI was applied to select a training set for model building and a test set for model validation. PCA was performed by Simca 14 (Umetrics, Sweden), whereas oCPLS2 and MLR-based analysis were implemented by in-house R-functions developed using R 4.0.4 (R Foundation for Statistical Computing, Vienna).

## 3 Results

PF were collected from corpses of both gender (with a M to F ratio of approximately 1:3), aged from 20 to 87 years (mean=51, SD=21) with a PMI ranging from 16 to 170 hours (mean=70, SD=46). Table 1 reports the demographic data, PMI, and causes of death. It is worth noting that PMI and age showed a moderate correlation (r=0.35, p=0.09), whereas no significant association was discovered for PMI and sex. A training set composed of 18 samples and a test set of 6 samples were selected. The distributions of sex and age in the training set were the same observed in the collected samples.

**Table 1.** Demographic parameters, PMI, and causes of death.

| sample | sex | age | PMI (h) | Cause of death |
| --- | --- | --- | --- | --- |
| 1 | F | 31 | 16 | Haemorrhagic shock |
| 2 | M | 20 | 20 | Mechanical asphyxia (hanging) |
| 3 | M | 35 | 25 | Acute Cardiac Failure |
| 4 | M | 21 | 32 | Epidural hematoma |
| 5 | M | 87 | 33 | Subdural hematoma |
| 6 | F | 52 | 33.5 | Cardiogenic shock (myocardial infarction) |
| 7 | M | 68 | 38 | Haemorrhagic/traumatic shock |
| 8 | M | 63 | 39 | Mechanical asphyxia (positional) |
| 9 | F | 81 | 41 | Acute Cardiac Failure |
| 10 | M | 32 | 46 | Drug intoxication (cocaine) |
| 11 | M | 38 | 46 | Mechanical asphyxia (hanging) |
| 12 | F | 60 | 62 | Pulmonary embolism and cerebellar ischemia |
| 13 | M | 42 | 63 | Cerebral ischemia |
| 14 | M | 37 | 66 | Haemorrhagic shock |
| 15 | M | 49 | 70.3 | Mechanical asphyxia (*ab ingestis*) |
| 16 | F | 39 | 77 | Multiorgan failure |
| 17 | M | 28 | 82 | Drug intoxication (cocaine, heroin) |
| 18 | M | 58 | 86 | Acute bowel infarction |
| 19 | M | 32 | 92 | Hypoxic brain death |
| 20 | M | 81 | 100 | Haemorrhagic shock |
| 21 | F | 77 | 120 | Neoplastic cachexia |
| 22 | M | 42 | 160 | Drug intoxication (buprenorphine) |
| 23 | F | 69 | 168 | Neoplastic cachexia |
| 24 | M | 80 | 170 | Acute Cardiac Failure |

Before NMR analysis, PF samples were treated in two different ways, namely ultrafiltration and liquid-liquid extraction, in view of the possibility of the future use of other analytical platforms based on Mass Spectrometry (MS) besides NMR. Indeed, MS techniques would allow us to expand the explorable metabolome of PF samples to other classes of molecules such as the more lipophilic ones. While NMR does not need strong manipulation of the biofluids, requiring only the elimination of large molecules such as proteins or lipids, MS techniques are mainly based on more invasive liquid-liquid extraction procedures with appropriate solvents. In this view, it is important to ensure that the extraction protocol does not significantly affect either qualitatively or quantitatively the metabolome. To this aim, we compared the spectra of ultrafiltered PF (UPF) with the spectra of the hydrophilic phase of the liquid-liquid extracted PF (LLEPF). We also recorded a spectrum of an untreated PF samples.

Fig.1 shows the aliphatic part of the ^1^H NMR spectrum of the untreated PF compared to the spectra of UPF and LLEPF. It is worth of note that the three samples were taken from the same individual, in order to compare the different experimental treatment without having to consider the interindividual variability. As can be seen, untreated PF (Fig 1A) exhibits broad resonances from macromolecules causing baseline distortion and interfering with the sharp resonances originating from low molecular weight metabolites. After removal of the macromolecules, both UPF (Fig. 1B) and LLEPF (Fig. 1C) give spectra with proper baselines showing only the required sharp resonances from small metabolites. Interestingly, the two NMR spectra of the protein-free samples are extremely similar. The main difference between them is the absence of ethanol in the LLEPF spectrum, as can be expected due to the sample concentration step required in the LLE procedure.

**Figure 1.**
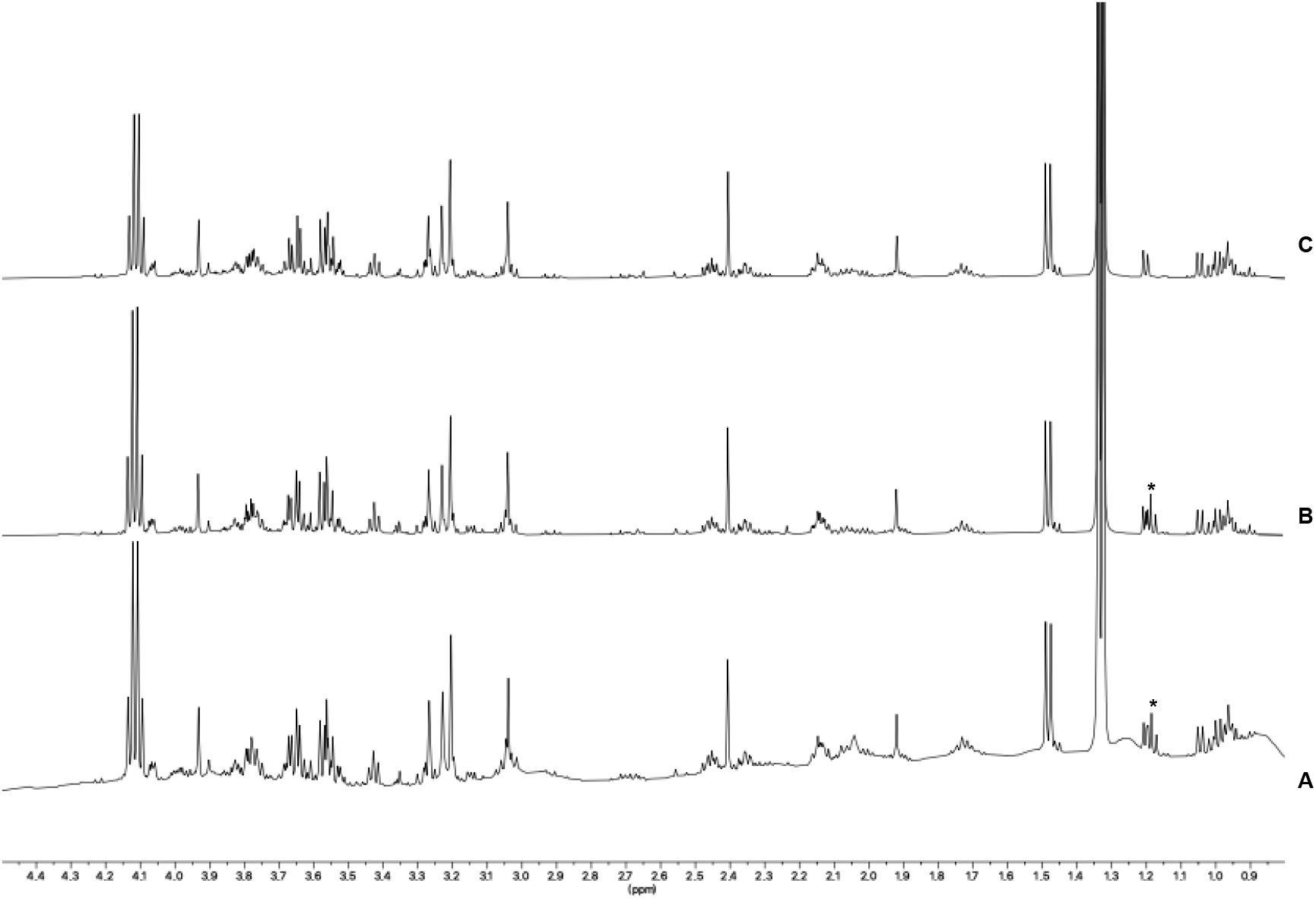
Aliphatic part of the ^1^H NMR spectrum of untreated PF, UPF and LLEPF samples. Three PF aliquots of the same individual were analysed. (*) indicates the triplet of ethanol.

24 PF samples collected during consecutive autopsies from cadavers with PMI ranging from 16 to 170 h were then processed using both experimental protocols. A total of 48 PF were analysed by ^1^H NMR. Comparison of all the 48 spectra recorded following the two experimental protocol, shows the absence of any significant qualitative or quantitative difference in the distribution of the recorded NMR resonances assigned to the small metabolites, indicating that proteins and/or other large molecules are equally excluded both with the physical procedure of filtering and with solvent extraction (Supplementary Fig. 1). 50 identified metabolites were quantified in both the datasets using the Chenomx NMR Suite Profiler tool (see Supplementary Table 1). The matrix of the quantified metabolites was submitted to multivariate analysis. Principal Component Analysis (PCA) was first applied to investigate trends and similarities among the spectra (Fig 2). No outliers were detected by T2 and Q-distance test assuming a significance level of 0.05 considering each single data set.

**Figure 2.**
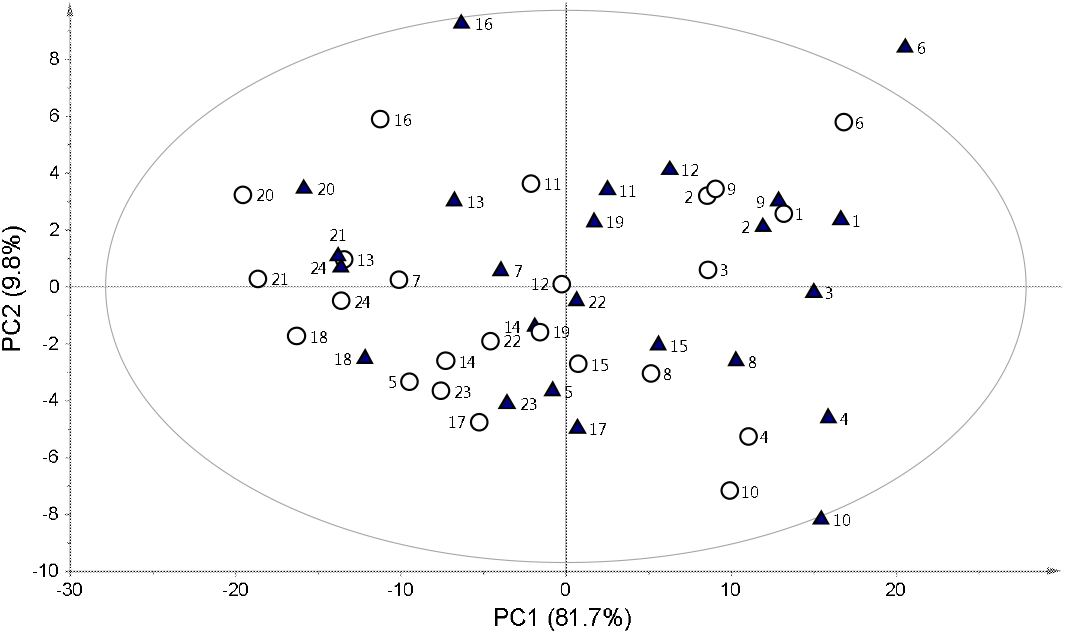
PCA model of PF samples. Samples are coloured accordind to the processing method (U: plain triangles; LLE: open circles) and labelled according to the individual.

As can be seen in Fig. 2, the two PF samples belonging to the same individual are very similar, lying in the same region of the multivariate space of the first two PCs, independently of the pre-analytical processing used. The observed scattering between the two samples of the same individuals reflects the sum of the experimental errors coming fron the extraction, the instrumental precision, the post-processing and fitting of the spectra, but it does not affect the global distribution of the samples and therefore the global metabolomic features.

Supervised data analysis based on Projection to Latent Structure regression (PLS) was applied to evaluate the effects of PMI on the metabolomic profiles of the collected PF samples. Considering the data set obtained from UPF samples, the oCPLS2 model showed 1 predictive score component, R^2^=0.547 (p=0.045), Q^2^=0.244 (p=0.010), standard deviation error in calculation (SDEC) equal to 30 hours, mean standard deviation error in cross-validatuon (SDECV) equal to 38 hours and standard deviation error in prediction (SDEP) equal to 33 hours. In Fig 3A, the PMI calculated for the training set and that predicted for the test set are reported. It is worth noting that samples with the highest PMI showed the worst estimate of PMI, with a negative deviation with respect to the experimental value. This may be due to a change of the biological mechanisms determining the metabolic content of PF at the highest PMI that may introduce a mild non-linearity in the behaviour of the metabolic concentration with respect to PMI. Unfortunately, the interval between 120 and 160 hours was not sampled and therefore it is not possible to investigate more in detail the behaviour of PMI in that interval, where non-linearity seems to appear. If PMI greater than 100 hours are not considered in the analysis, a oCPLS2 model with 1 predictive score component, R^2^=0.543 (p=0.030), Q^2^=0.444 (p=0.005), SDEC=17 hours, SDECV=19 hours and SDEP=15 hours, can be obtained. As it can be observed in Fig. 4A, where PMI calculated for the training set and that predicted for the test set are reported, the behaviour of PMI estimated by the model, i.e. the predictive score component, shows a linear behaviour with respect to the experimental PMI. Stability selection applied to the model built considering the whole range of PMI discovered 17 metabolites as significantly relevant, 10 of which resulted to be significantly releated to PMI after correction by age (Table 2).

**Figure 3.**
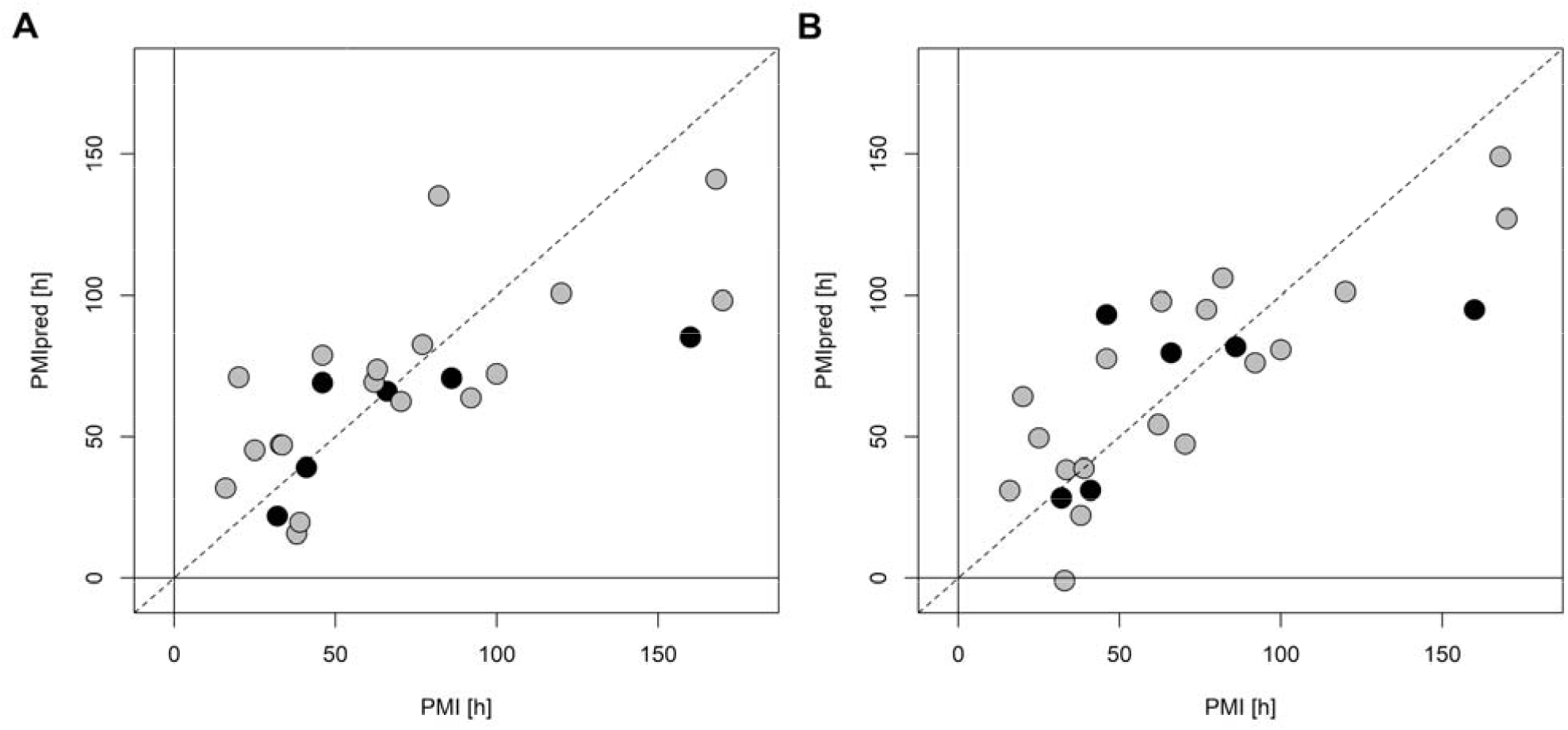
PMI calculated for the training set (grey circles) and predicted for the test set (black circles) vs experimental PMI for in the range [16,170] h. Panel A reports the results for the data obtained from UPS samples, while panel B those from LLEPF.

**Figure 4.**
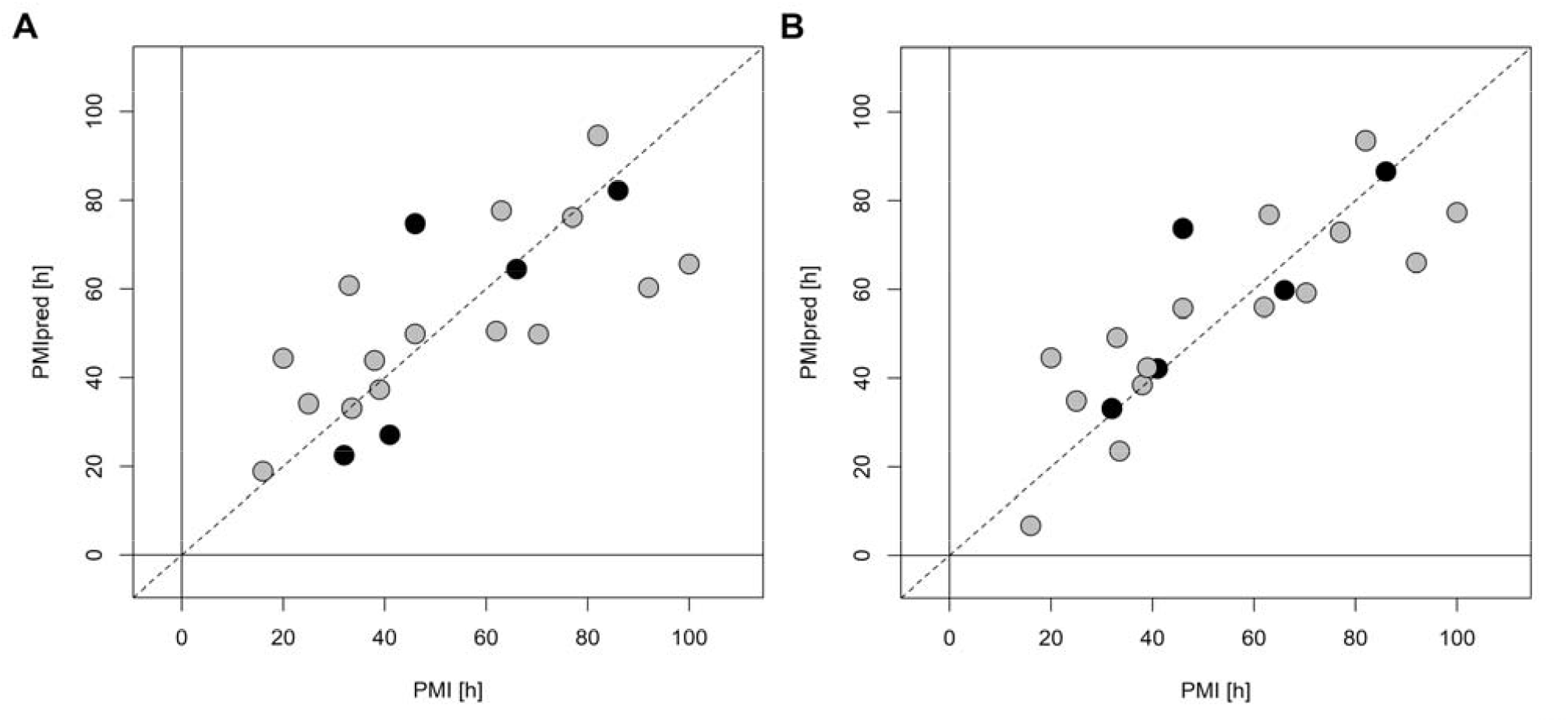
PMI calculated for the training set (grey circles) and predicted for the test set (black circles) vs experimental PMI for in the range [16,100] h. Panel A reports the results for the data obtained from UPS samples, while panel B those from LLEPF.

**Table 2.** Relevant metabolites discovered by oCPLS2 analysis; protocol specifies the type of experimental protocol applied for protein removal, r is the Pearson’s correlation coefficient between metabolite concentration corrected by age and PMI, and p is the p-value of the coefficient of PMI in the MLR regression model that uses PMI and age as factors and metabolite concentration as response.

| Metabolite | Protocol | r | p |
| --- | --- | --- | --- |
| Acetate | LLE | 0.387 | 0.169 |
|  | U | 0.403 | 0.150 |
| Betaine | LLE | 0.199 | 0.495 |
|  | U | 0.317 | 0.267 |
| Butyrate | U | 0.641 | **0.012** |
| Choline | LLE | 0.539 | **0.043** |
|  | U | 0.606 | **0.019** |
| Citrate | LLE | -0.605 | **0.019** |
| Ethanolamine | LLE | 0.633 | **0.013** |
|  | U | 0.632 | **0.013** |
| Formate | LLE | 0.554 | **0.037** |
|  | U | 0.556 | **0.036** |
| Glycine | LLE | 0.728 | **0.002** |
|  | U | 0.723 | **0.003** |
| Hypoxanthine | LLE | -0.630 | **0.014** |
|  | U | -0.559 | **0.035** |
| Inosine | LLE | -0.419 | 0.132 |
|  | U | -0.358 | 0.206 |
| Mannose | LLE | -0.436 | 0.115 |
| Methanol | LLE | -0.461 | 0.093 |
|  | U | -0.267 | 0.355 |
| Myo-inositol | U | -0.036 | 0.902 |
| Nicotinurate | LLE | 0.255 | 0.377 |
| Proline | U | 0.641 | **0.011** |
| Propionate | U | 0.536 | **0.045** |
| β-Alanine | U | 0.279 | 0.333 |
| Trimethylamine | LLE | 0.508 | **0.060** |
|  | U | 0.561 | **0.034** |
| Uracil | U | 0.700 | **0.004** |
| Uridine | U | 0.266 | 0.357 |

The data set obtained from LLEPF samples led to a regression model with 1 predictive and 1 non-predictive score component, R^2^=0.669 (p=0.065), Q^2^=0.161 (p=0.045), SDEC=25 hours, SDECV=41 hours, and SDEP=34 hours. In Fig 3B, the PMI calculated for the training set and that predicted for the test set are reported. Also in this case, a mild non-linearity is observed in the data. If PMI greater than 100 hours are not considered, a oCPLS2 model with 1 predictive score component, R^2^=0.701 (p=0.005), Q^2^=0.569 (p=0.005), SDEC=14 hours, SDECV=17 hours and SDEP=13 hours, can be obtained. The PMI calculated for the training set and that predicted for the test set are reported in Fig. 4B. As in the case of the data set obtained by ultrafiltration, the behaviour of PMI estimated by the model shows a linear behaviour with respect to the experimental PMI. Applying stability selection to the model built considering the whole data set, 13 metabolites resulted to be relevant, 7 of which resulted to be significantly releated to PMI (Table 2).

The profiles of the relevant metabolites mainly related to PMI common to both data sets are reported in Fig. 5 (data from UPF samples are shown). Similar profiles were observed for the data obtained from LLEPF samples (data not shown). Interestingly, formate and trimethylamine show a sudden increase at the highest PMI. The same trend is observed for propionate in UPF samples. This may be due to the presence of outliers or to a real change in the biological mechanism underlying PMI. Further investigations are requested to inverstigate more in depth this finding. When the reduced PMI range (16-100 hours) is considered, the relevant metabolites related to PMI do not change except for formate and trimethylamine that are not significant anymore, further corroborating the hypothesis that they are only related to higher PMIs.

**Figure 5.**
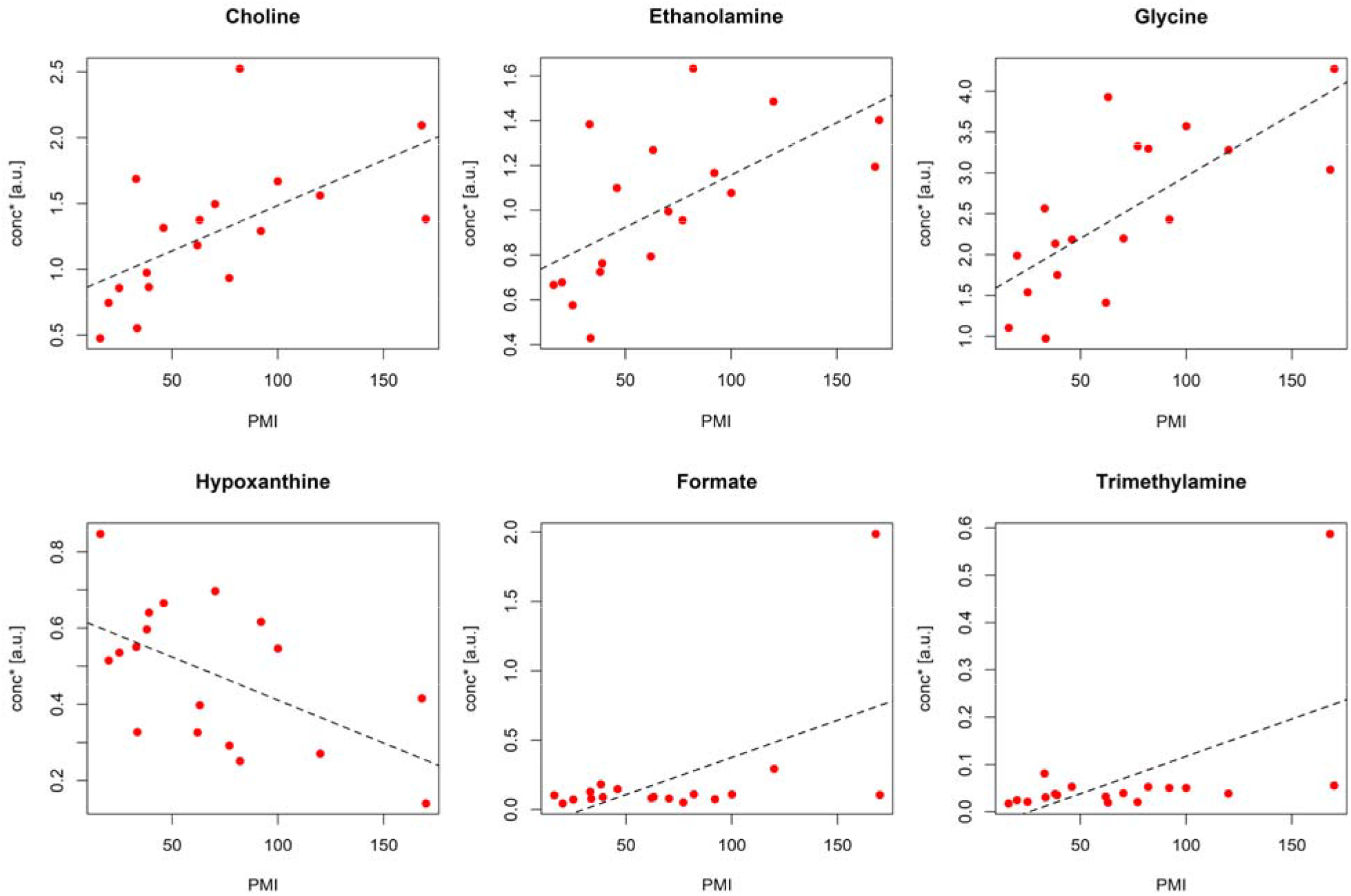
Data obtained from UPF dataset: profiles of the metabolite concentration corrected by age (conc*) vs PMI for the relevant metabolites discovered using datasets from both UPF and LLEPF. Only metabolites with p less than 0.10 are reported.

## 4. Discussion

Considering that little is known about the PF metabolomic composition, our first objective was to establish whether the analytical protocol used to extract the metabolites could influence the detectable metabolome. PCA analysis clearly shows that the distribution of PF samples is maintained regardless of the protocol used, indicating that the explored metabolome remains unaffected. In our previous works, we investigated post-mortem metabolomics through ^1^H NMR analysis of biofluids processed via ultrafiltration [17, 18, 20, 21]. However, in the perspective of using multiple analytical platforms, expanding toward mass spectrometry, we explored the use of two different experimental protocols. The main difference we observed is that LLE allows to rule out ethanol, typically present in high concentrations in real-life forensic samples, the NMR resonances of which is overlapped to signals of endogenous molecules of interest (among all 3-hydroxybutyrate).

Furthermore, since sampling was essentially performed with no exclusion criteria, except for obvious PF pathological appearance, the metabolomic composition did not appear related to sex and cause of death, although a moderate correlation was observed between PMI and age.

Post-mortem modifications in the PF metabolome were found to be correlated with PMI, so that a predictive model for PM estimation was built and rigorously validated with an independent test set of PF samples. The error in prediction was very similar in the two experimental datasets (33 and 34 hours for U and LLE, respectively) over the studied PMI window (16-170 hours). Samples with PMI greater than 100 hours were characterised by the highest errors in prediction, suggesting an underlying biological mechanism which could result in a non-linear behaviour. This is somehow expectable as bacterial growth occurs at later PMIs, resulting in a co-metabolism. We have previously observed this phenomenon in a highly controlled animal model of PMI estimation on aqueous and vitreous humour samples [17, 18, 20]. Since in the present experiment the interval between 120 and 160 hours, when non-linearity seems to appear, was not adequately sampled, it was not possible to better investigate the behaviour of the metabolome in this interval. Interestingly, some of the longer PMI samples display however several features proper of bacterial metabolism, namely formate, trimethylamine, propionate and butyrate.

Due to the aforementioned, we have decided to focus on the more homogeneously covered PMI window, i.e., up to 100 hours. This strategy led to a remarkable improvement of the prediction model in both experimental protocols with errors in prediction of 15 and 13 hours, for U and LLE, respectively.

The design of the experiment was certainly underpowered to make any conclusive inferences about the underlying metabolic phenomena. Despite this, 20 metabolites were relevant in the PMI regression, 11 out of 20 were shared by U and LLE protocols. Among the latter, choline, ethanolamine, glycine, hypoxanthine, formate and trimethylamine were statistically significant (p <0.10) with respect to PMI. Taking into consideration PMI up to 100 hours, formate and trimethylamine were no longer significant, confirming their role only in very late PMI. Of note, among the 4 remaining metabolites, hypoxanthine was the only one with a decreasing behaviour, whereas choline, ethanolamine and glycine were directly related to PMI.

Mizutani et al. observed stable 3-hydroxybutyrate levels in post-mortem human PF within 96 hours after death [12]. Interestingly, this trend was also observed in our data, even in samples with higher PMIs (up to 170 hours), confirming this metabolite’s resilience to post-mortem in human PF.

Our study has several limitations. Firstly, due to its proof-of-concept nature, only a limited number of samples were analysed. Secondly, random sampling did not allow to homogeneously cover the entire time-window investigated, with an underrepresentation of PMI >100 hours. Thirdly, PFs were taken from consecutive judicial cases and in many of them the PMI was estimated through circumstantial evidence and/or classical thanatological signs; most of the corpses were stored cooled in the morgue before autopsy for an uneven time. Finally, the samples were analysed through a single analytical platform which is not appropriate for investigating the lipophilic phase.

Despite these limitations, the presented results show that the spectrum of biological activity associated with post-mortem modifications in a real-life scenario can be intercepted by the proposed approach, independently of a number of major variables, namely cause and mechanism of death, sex and corpse storage. Additionally, notwithstanding the limited sampling, PMI was accurately predicted using an independent set of samples with error as low as 13 hours for PMI ≤100 hours. Moreover, we demonstrated that the metabolite extraction procedure did not alter the metabolome composition and can therefore be independently selected according to the chosen analytical platform.

## 5. Conclusions

In conclusion, PF was found to be a biofluid of interest for post-mortem metabolomics. Hence, this proof-of-concept approach deserves to be translated on a wide dataset and investigated with different analytical platforms. Due to the peculiar physiopathology, PF metabolomics appears intriguing to investigate the cause and mechanism of death, unless modifications related to PMI are fully taken into account, either standardising the samples according to PMI or constraining the model to this factor.

## Declarations

## Authors’ contributions

All authors read and approved the final manuscript. A.C. and M.S. equally contributed to this work.

## Data availability

Datasets generated and/or analysed during the current study are available from the corresponding author on reasonable request.

## Ethics declarations

As samples were obtained during judicial autopsies informed consent was obtined through local Prosecutor Office.

