## Supplementary material for "Metabolomics investigation of post-mortem human pericardial fluid": Supplementary Material.docx

**Table S1.** List of human PF metabolites quantified using the Chenomx Profiler tool.

| **Compound** | **PubChem (CID)** |
| --- | --- |
| 3-Hydroxybutyrate | 92135 |
| Acetate | 176 |
| Acetone | 180 |
| Alanine | 5950 |
| Asparagine | 6267 |
| Aspartate | 5960 |
| Betaine | 248 |
| Butyrate | 264 |
| Choline | 305 |
| Citrate | 311 |
| Creatine | 586 |
| Creatinine | 588 |
| Dimethylamine | 674 |
| Ethanolamine | 700 |
| Formate | 284 |
| Fumarate | 723 |
| Glucose | 5793 |
| Glutamate | 33032 |
| Glutamine | 5961 |
| Glycerol | 753 |
| Glycine | 750 |
| Histidine | 6274 |
| Hypoxanthine | 790 |
| Inosine | 6021 |
| Isoleucine | 6306 |
| Lactate | 108689 |
| Leucine | 6106 |
| Lysine | 5962 |
| Maltose | 439186 |
| Mannose | 18950 |
| Methanol | 887 |
| Methionine | 6137 |
| Nicotinurate | 68499 |
| Ornithine | 6262 |
| Phenylalanine | 6140 |
| Proline | 145742 |
| Propionate | 1032 |
| Serine | 5951 |
| Succinate | 1110 |
| Taurine | 1123 |
| Threonine | 6288 |
| Trimethylamine | 1146 |
| Tryptophan | 6305 |
| Tyrosine | 6057 |
| Uracil | 1174 |
| Uridine | 6029 |
| Valine | 6287 |
| myo-Inositol | 892 |
| sn-Glycero-3-phosphocholine | 439285 |
| β-Alanine | 239 |

**Fig. S1**. ^1^H NMR spectra of (A) UPF and (B) LLEPF samples. Three PF aliquots of the same individual were analysed. (*) indicates the resonances of ethanol.

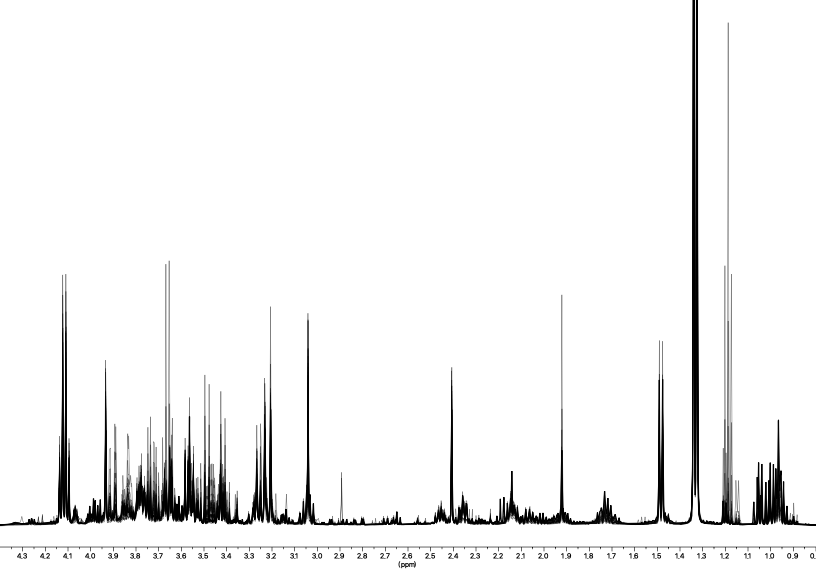

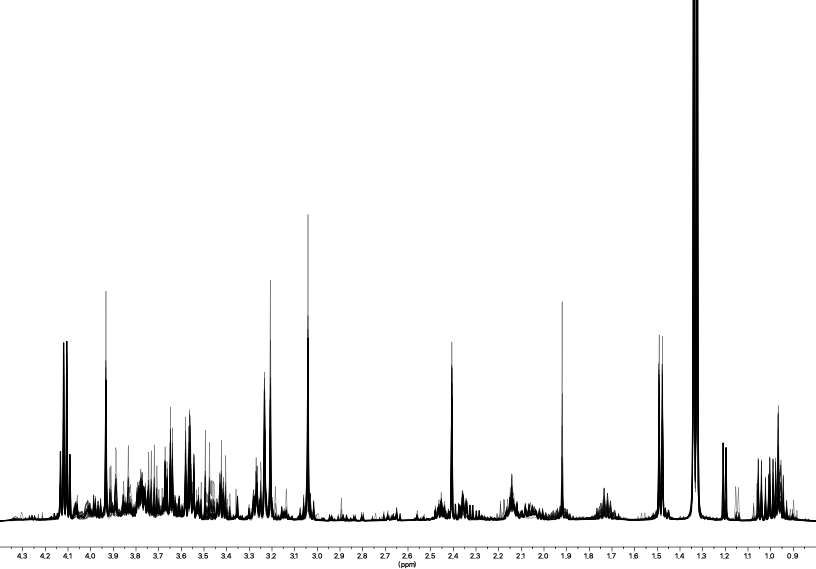

*****

*****

**B**

**A**
